# Contrasting needle physiological strategies to soil nutrient scarcity in radiata pine plantations

**DOI:** 10.1101/2025.07.15.664982

**Authors:** Lorena Ruiz de Larrinaga, Francisco San Miguel-Oti, Ander Monasterio, Unai Artetxe, Sergi Munné-Bosch, Tania Mesa, Celine Moreaux, Unai Ortega-Barrueta, Unai Sertutxa, Lorena Peña, Ibone Ametzaga-Arregi, Jorge Curiel Yuste, Raquel Esteban

## Abstract

Intensive European forestry practices contribute to soil degradation and nutrient depletion, compromising tree health and ecosystem stability. However, the influence of soil nutrient scarcity on tree physiological responses during drought remains unclear, particularly in *Pinus radiata* D. Don plantations, where it may impair needle function and overall tree health. We investigated how soil nutrient availability influences leaf-level physiological strategies during drought by comparing same-aged needles from two *P. radiata* stands with contrasting management stages and ages: (i) trees (≈20 years), likely to be clear-cut at 30-35 years (herein managed), and (ii) trees (>45 years) not clear-cut at the typical rotation age (herein abandoned). During the severe summer drought of 2022, both stands exhibited downregulation of the photosynthetic apparatus, indicating impaired photosynthetic performance under drought. However, their leaf-physiological responses to nutrient scarcity diverged. Managed trees exhibited dependence on soil nutrients, with reduced photosynthetic performance under nutrient-poor conditions. In contrast, abandoned trees showed relative independence from soil nutrients, maintaining photosynthetic function even under nutrient-poor conditions. This highlights a trade-off: younger, managed trees may enhance photosynthesis under optimal nutrient conditions but are more susceptible to nutrient imbalances and environmental stress. In turn, older, abandoned trees appeared to buffer the effects of drought and nutrient scarcity through age-related physiological traits. Our findings underscore the physiological value of mature stands. Furthermore, higher soil organic carbon and vegetation diversity in abandoned stands suggest that management cessation may enhance long-term ecosystem resilience. These results emphasise the importance of integrating soil–leaf interactions into sustainable forest management.

## Introduction

Global wood demand is forecasted to double, increasing from 3.4 billion m^3^ in 2010 to 7.6 billion m^3^ by 2030 (UNECE/FAO 2021). To meet this growing demand, many regions in Europe have adopted a forestry model based on increasingly intensive management practices, such as rotations (35–40 years rotations) and clear-cutting(McGrath et al., 2015). This model boosts short-term timber production but, on the other hand, compromises essential ecosystem functions such as nutrient cycling, soil stability, and biodiversity conservation (Sing et al. 2018; Sertutxa et al. 2024). Natural disturbances such as droughts put further pressure on forests by increasing tree mortality and reducing productivity (Senf et al., 2020). In the northern Iberian Peninsula, the spread of fast-growing evergreen *P. radiata* D. Don plantations during the 20th century significantly increased timber production but also introduced problems such as soil compaction, erosion, and nutrient depletion due to the type of management implemented (Merino et al., 2004). The loss of nutrients and disruptions or imbalances in nutrient cycling are strongly linked to reduced forest productivity and ecosystem stability (Jonard et al., 2015). This ongoing nutrient depletion could threaten the long-term viability of forests, putting at risk their ability to meet the rising global demand for wood.

The availability of key soil nutrients, especially nitrogen, phosphorus, and potassium (i.e., NPK), is often reflected in the nutrient content of tree foliage(Davis et al., 2007). As a result, analysing foliar nutrients is a useful way to detect imbalances, which can reveal soil degradation and support forest health monitoring (ICP Forest; Michel et al. 2023). However, the relationship between soil and plant nutrients is complex. Plants regulate nutrient uptake based on their physiological needs and soil nutrient availability (Smith-Ramesh & Reynolds, 2017). Depending on their developmental stage (Yuan and Chen 2010; Chen et al. 2016), overall health (Morillas et al., 2012), soil properties (Kou et al., 2017) and environmental factors (e.g., elevation, temperature, precipitation, etc.) (Wang et al. 2022; Zhang et al. 2018;Yan et al. 2018), plants may prioritise nutrient conservation and remobilisation (allocation strategy) over nutrient acquisition. This dynamic response to optimise resource use underscores the plant’s ability to balance growth with acclimation.

Changes in nutrient availability and allocation can significantly influence the functionality of photosynthetic apparatus and overall plant physiological traits (Kumar et al., 2021), including nutrient-dependent regulation of phytohormones (Jia et al., 2022). Phytohormones are recognised as central regulators of plant growth and development (Davies, 1987) and orchestrate plant performance at the whole-plant level (Müller & Munné-Bosch, 2021), influencing not only life cycle processes but also the plant’s ability to respond to various environmental stresses. Among the phytohormones catalogued as stress hormones, abscisic acid (ABA) is a key player, modulating critical physiological processes such as stomatal closure, while enabling plants to cope with both abiotic and biotic stresses (Kundu & Gantait, 2017). Jasmonates, including jasmonic acid (JA), its bioactive derivative jasmonoyl-isoleucine (JA-Ile), and its biosynthetic precursor 12-oxo-phytodienoic acid (OPDA), act as major regulators of plant defence mechanisms and stomatal regulators together with ABA (Hewedy et al. 2023;Savchenko et al. 2014). Salicylic acid (SA) further contributes to the plant’s defence mechanism, playing a critical role in pathogen resistance, abiotic stress tolerance (together with JA and ABA), and systematic acquired resistance (Ghosh & Roychoudhury, 2024). Additionally, ethylene is essential for many developmental processes, and it is considered to play a major role as a signal molecule at low concentrations in the tolerance of several species to biotic and abiotic stresses (Müller & Munné-Bosch, 2015). Ultimately, melatonin (Mel) (N-acetyl-5-metroxytryptamine), an indole derivative (Dubbels et al., 1995), plays an important role in multiple physiological responses as a possible key regulator in plants (Arnao and Hernández-Ruiz 2015, 2018, 2021; Arnao et al. 2022). Mel contributes to every stage in the sequence of events triggered by an environmental stimulus, increasing tolerance against a stressor (Khan et al. 2020;Arnao et al. 2022). An extensive analysis of current studies shows that Mel regulates the metabolism, signalling and response of plant hormones such as auxin, gibberellins, cytokinins, ABA, ethylene, SA, jasmonates, brassinosteroids, polyamines, and strigolactones (Arnao & Hernández-Ruiz, 2021). Among physiological processes, Mel improves the rate and efficiency of photosynthesis (Yang et al., 2022), and regulates the metabolism of carbohydrates, lipids, and nitrogenous and sulfur compounds (Erdal, 2019). Additionally, Mel contributes to mineral ion homeostasis and supports nutrient acquisition under stressful conditions or nutrient deficiencies. Specifically, evidence indicates that Mel promotes nitrogen uptake, sustains potassium homeostasis, and plays a role in mitigating phosphorus deficiency (Sun et al., 2022).

Phytohormones are crucial in coordinating plant responses to environmental challenges, particularly by modulating photoprotection mechanisms. Among these, carotenoids are essential for fine-tuning plant-environment interactions, serving as antioxidants and energy dissipators to mitigate damage from environmental stresses (Gómez-Sagasti et al., 2023). Indeed, the interplay between photoprotective isoprenoids, the dynamic xanthophyll cycle (VAZ) (involving the three carotenoids, violaxanthin (V), antheraxanthin (A) and zeaxanthin (Z)), together with tocopherols (Munné-Bosch, 2005), contributes to safeguarding the photosynthetic apparatus from oxidative stress.

The crosstalk between plant performance (mediated by phytohormones, photoprotective compounds and photosynthetic efficiency) and nutrients is highly complex. This interplay involves multiple signalling pathways, which ensure finely tuned acclimation to the environmental conditions (Müller and Munné-Bosch 2021; Ahmad et al. 2023). Despite its importance, current knowledge remains limited in the following two key areas: i) the relationship between phytohormones and nutritional homeostasis, particularly in response to soil NPK availability, and ii) the role of phytohormonal regulation in plant physiological processes, such as photochemical efficiency and photoprotective mechanisms, in long-lived organisms like trees. Addressing these complexities requires integrating physiological insights with empirical data from tree plantations and management practices to identify key relationships that drive resilience. Understanding the intricate interplay between nutrients (both in soil and leaves), phytohormones, and photoprotective isoprenoids is essential. Such an approach is crucial not only for advancing fundamental knowledge of plant biology but also for its significant implications in fostering a sustainable bio-based economy.

Therefore, in this paper, we aimed to explore the physiological responses to a severe dry summer in 2022 in same-aged needles of two *P. radiata* stands, differentiated by their management stages and stand ages: (i) >45-year-old pine trees where management was abandoned (herein abandoned) and (ii) ≈20-year-old managed pine trees subjected to thinning, pruning, and clearing (herein managed). Importantly, although the stands differ in age, all sampled needles were of the same age cohort, allowing a direct comparison of leaf-level physiological traits. We therefore hypothesised that under contrasting management stages in *P. radiata* stands, and hence under contrasting nutrient status, trees will adjust their physiological performance by regulating the relationship between soil nutrient availability and leaf nutrient status, resulting in distinct strategies to maintain photosynthetic function. For managed, younger trees, we expected photosynthesis to have a stronger dependence on soil nutrient availability, with phytohormones playing a critical role in modulating leaf nutrient status and physiological performance. The reliance on soil nutrients would drive a more acquisitive physiological strategy, prioritising photosynthesis and vitality at the cost of resilience. Conversely, abandoned, mature trees would adopt a conservative physiological strategy, utilising nutrient remobilisation to maintain photosynthetic function independently of external nutrient availability. This research will provide new insights into the complex interactions between soil conditions, leaf nutrient status, stress hormones, and plant physiological responses under varying management stages and ages, contributing to the development of more sustainable forestry strategies in the face of climate change and escalating global wood demand.

## Materials and Methods

### Study area description

The study was conducted in July 2022 in *P. radiata* D. Don stands of the Urdaibai Biosphere Reserve (UBR), a site of significant ecological value located on the coast of the Bay of Biscay in the Basque Country, Spain (43°19′ N, 2°40′ W). In the UBR, the surge in demand for wood driven by industrialisation in the 1950s led to the abandonment of crops and meadows (>40 years) and the spread of fast-growing *P. radiata* plantations, resulting in significant fragmentation of native forests. Thus, the native vegetation comprises Atlantic mixed forests that occupy 9% of the territory, while forest plantations of exotic species (*P. radiata* and *Eucalyptus spp.*) occupy 50% of the area (Castillo-Eguskitza et al., 2017), which are characterised by acid soils with a pH between 3.5 and 6 (Loidi et al., 2011). The climate in the area is temperate Atlantic, with a mean monthly temperature and total monthly rainfall ranging from 7°C and 64.1 mm in January, to 21.8°C and 12 mm in July 2022 (CRU TS v.4.07; Harris et al. 2020, reference period 2022) while the mean Standardised Precipitation-Evapotranspiration Index for 2022 (Mean SPEI_2022_) was – 1.33 ± 0.88 (Global SPEI database; (Vicente-Serrano et al. 2010; Beguería et al. 2010, 2014). SPEI is a multiscale drought index, based on temperature and evapotranspiration data, that indicates drought conditions when registering negative values. The SPEI_2022_ value thus indicates severe drought conditions for the study year (Spinoni et al. 2019; García-Valdecasas Ojeda et al. 2021).

### Radiata pine plantations selection

Two types of *P. radiata* plantations within the UBR were considered to analyse how management and stand age influence the relationship between nutrients (both in the soil and needles), and their impact on tree physiological functioning: (i) > 45 years pine stands characterized by trees that have not been clear-cut at the rotation age (30-35 years), whose management has ceased (herein abandoned) and (i) managed pine stands (∼22 years trees) subjected to thinning, pruning and clearing (herein managed) that will be clear-cut when reaching 30-35 years. For each type, four stands were selected, all with similar characteristics (i.e. soil properties, topography, stand density, historical land use, diseases, climatic conditions, tree age) despite not being close to one another. For each stand, a central point (centroid) was established randomly, taking care to ensure homogeneous characteristics. In total, eight centroids were sampled (Figure 1A). To achieve a representative sample of both types of management practices, orthophotos were used to confirm the developmental stage of managed stands and to ensure the abandonment stage (>45 years) of abandoned ones. Also, field visits were conducted to ensure the absence of recent management activities in abandoned areas.

**Figure 1.**
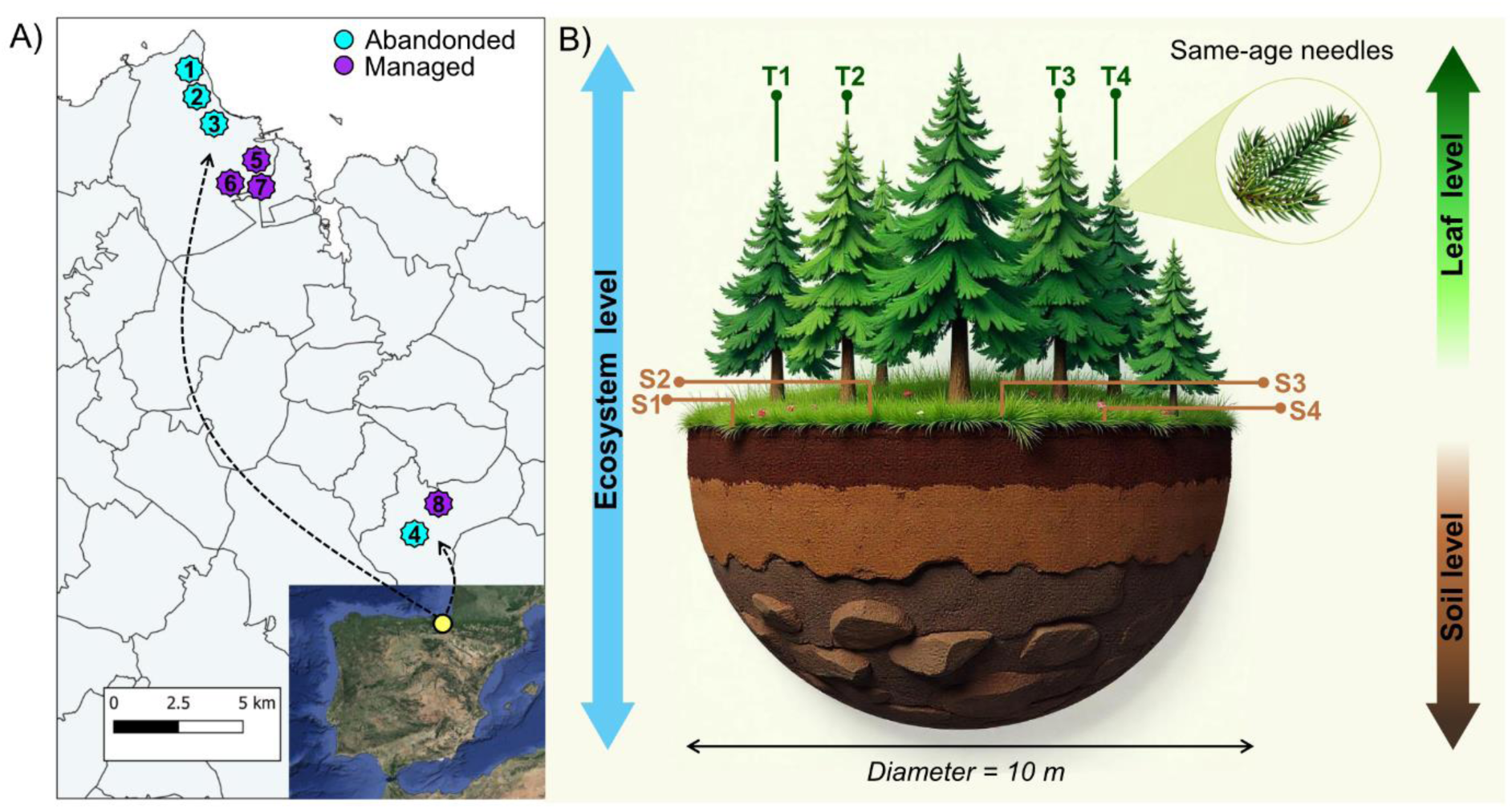
A) Geographical distribution of the eight study centroids (four centroids per condition) within the northern area of the Iberian Peninsula (abandoned and managed forest, marked in blue and purple, respectively). B) In each centroid of 10 metres diameter, four adult *P. radiata* trees were randomly selected for a total of 16 individuals per forestry practice to measure variables at ecosystem, soil and tree levels. The soil was sampled from four random points across each centroid (S_1_, S_2_, S_3_, S_4_) and 2- to 3-year-old mature needles were sampled randomly from the four cardinal directions within each tree (T_1_, T_2_, T_3_, T_4_).

### Sampling design

In each centroid, soil samples were taken at 20 cm depth from four random points across each centroid to account for the soil nutrient heterogeneity. Once collected, soil samples of each centroid were mixed and immediately stored in a portable fridge and maintained at 4°C until they were transported to the laboratory, where samples were dried at room temperature (i.e. 20°C), sieved using a 2 mm mesh size and stored in darkness for further analyses. Also, the four closest trees were randomly selected, resulting in a total of 16 trees across the four centroids. To characterise them based on their physiological traits (i.e. tree characterisation, see below), data were collected from a total of 32 trees (2 types of management practices x 4 centroids per management practice x 4 trees per centroid) (Figure 1B). The selected trees showed no evident signs of health loss. The measured mean diameter at breast height (DBH, i.e., the trunk diameter 1.3 m above the ground) was 54.83 ± 1.66 cm and 35.84 ± 1.53 cm for abandoned and managed trees, respectively (Table 1).

**Table 1.**
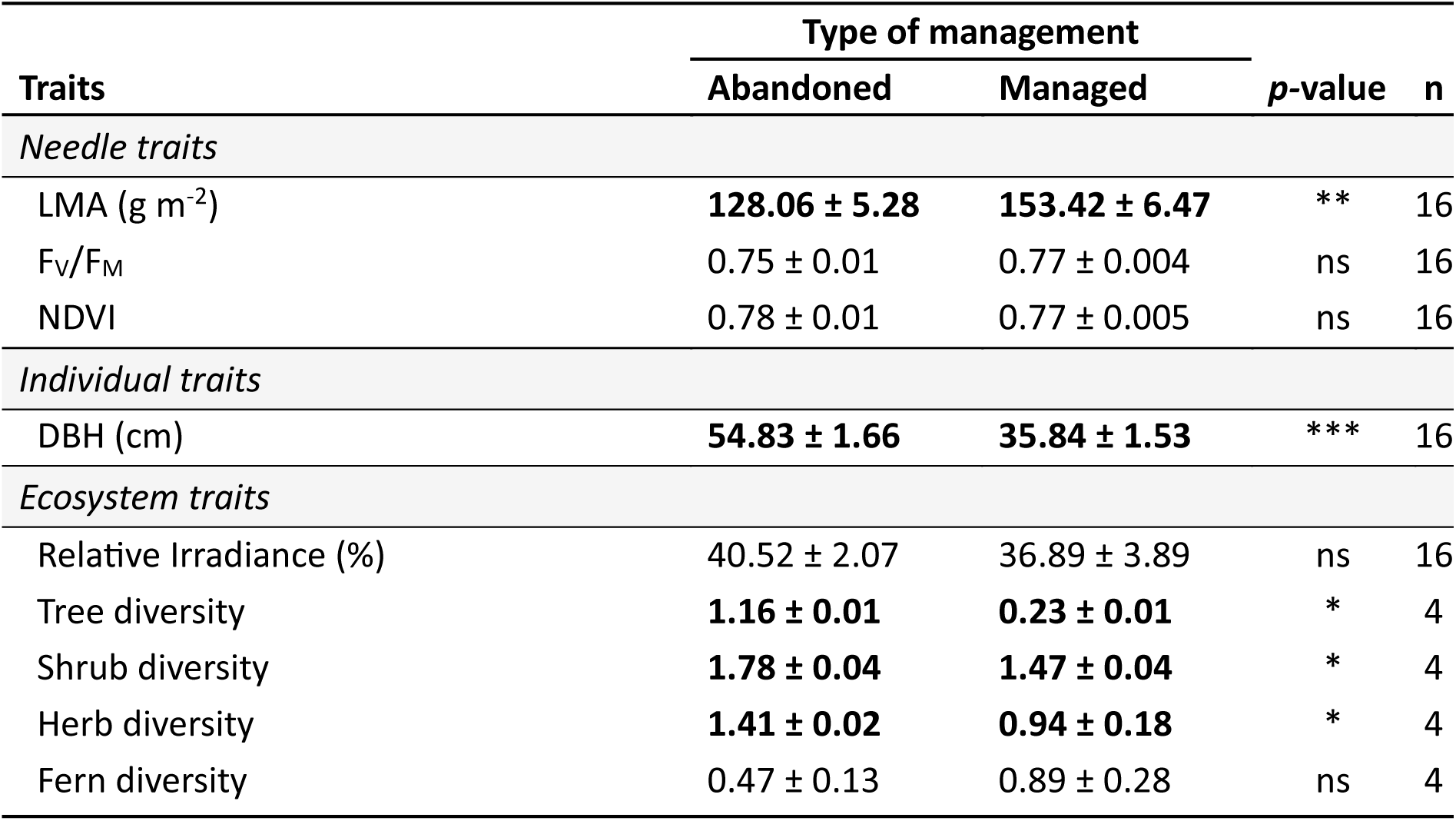
Needle, individual tree and ecosystem traits between trees growing in two types of management practices, labelled as abandoned and managed. (i.) individual tree traits as leaf mass per area (LMA), photochemical efficiency (F_V_/F_M_), leaf-level normalised difference vegetation index (NDVI) and diameter at breast height (DBH). (ii.) Ecosystem traits, including relative irradiance and tree, shrub, herb and fern diversity, are shown. Values are the mean of 16 replicates ± standard error. Additionally, the table includes p-values after one-way ANOVA statistical analysis, with asterisks indicating the level of statistical significance between the abandoned and managed groups: * for p < 0.05, ** for p < 0.01, and *** for p < 0.001. When normality and homocedasticity assumptions were not met, the Kruskal-Wallis test was used.

| Traits | Type of management |  | *p*-value | n |
| --- | --- | --- | --- | --- |
|  | Abandoned | Managed |  |  |
| *Needle traits* |  |  |  |  |
| LMA (g m<sup>-2</sup>) | 128.06 ± 5.28 | 153.42 ± 6.47 | ** | 16 |
| F<sub>V</sub>/F<sub>M</sub> | 0.75 ± 0.01 | 0.77 ± 0.004 | ns | 16 |
| NDVI | 0.78 ± 0.01 | 0.77 ± 0.005 | ns | 16 |
| *Individual traits* |  |  |  |  |
| DBH (cm) | 54.83 ± 1.66 | 35.84 ± 1.53 | *** | 16 |
| *Ecosystem traits* |  |  |  |  |
| Relative Irradiance (%) | 40.52 ± 2.07 | 36.89 ± 3.89 | ns | 16 |
| Tree diversity | 1.16 ± 0.01 | 0.23 ± 0.01 | * | 4 |
| Shrub diversity | 1.78 ± 0.04 | 1.47 ± 0.04 | * | 4 |
| Herb diversity | 1.41 ± 0.02 | 0.94 ± 0.18 | * | 4 |
| Fern diversity | 0.47 ± 0.13 | 0.89 ± 0.28 | ns | 4 |

For each tree, two to three-year-old mature needles (*P. radiata* may retain their leaves for up to four years) were sampled randomly with a pole saw from the four cardinal directions of each tree. Despite the difference in tree ages, our study focuses exclusively on needles of the same age in both management stages, allowing for direct physiological comparisons across stands. Since leaf compounds (i.e., pigments and tocopherols) exhibit a high degree of environment modulation(Esteban et al., 2015), needle samples were immediately stored in hermetic plastic bags where relative humidity, temperature, and darkness conditions were maintained constant for about 12 h. This pretreatment was meant to avoid circadian effects, thus facilitating the comparison among different sampling study sites and days(Fernández-Marín et al., 2019). Due to logistics (i.e., the number of sites and individuals in each site in a wide sampling area), sampling was performed on different days and at different hours. Upon completing the pretreatment, the photochemical efficiency and chlorophyll *a* fluorescence induction (OJIP) were assessed. For hormone, pigment, and tocopherol analysis (see details below), needle samples were immediately placed in liquid nitrogen and stored at -80°C. For leaf nutrient analysis, needle samples were dried at 80°C for 48 h, ground in a blender (IKA Multidrive Control, Staufen, Germany), and stored at room temperature with silica gel (to avoid tissue rehydration) until analysis.

### Ecosystem traits characterisation

Each forest stand was characterised by the following traits (Table 1): relative irradiance, tree diversity, shrub diversity, and herbaceous diversity. The relative irradiance expressed as a percentage (%) is defined as the quantity of solar radiation below each selected pine tree (Jennings, 1999). To calculate it, canopy photos with a digital camera (Nikon Coolpix 4500, Japan) equipped with a Nikon Fisheye Converter and mounted on a tripod for levelling and stabilisation, were taken with a hemispherical lens (for later environmental light determination). All hemispheric photographs were taken facing south, 50 cm from the tree trunks, and 2 m above the ground and were analysed by using Gap Light Analyser software (version 2.0; Frazer et al. 1999).

The diversity of each life form was analysed in each centroid by selecting nine plots of 10 m^2^ (5x2 m) separated by a distance of 5 m to identify and classify plant species into four groups according to their life form (trees, shrubs, herbs, and ferns) as performed by (Sertutxa et al., 2024). The diversity of each life form was calculated using the Shannon diversity index (Shannon and Weaver 1949).

### Individual tree traits characterisation

For the tree characterisation, the following traits were calculated (Table 1): leaf mass per area (LMA), leaf level-normalised difference vegetation index (NDVI), and OJIP (see below). In detail, for LMA calculation (expressed as grams of dry leaf per needle area in m^2^), photographs of three random needles from each individual were taken with a digital camera (Nikon Coolpix 4500, Japan) and processed with the Java-based image program Image J (1.52p; Ferreira and Rasband 2019) to quantify the area of the needle. Immediately, these same needles were weighed to obtain fresh weight, dried at 80°C for 48 h, and weighed again to estimate dry weight. The NDVI was measured in the adaxial side of needles using a handheld spectrophotometer (SpectraPen SP 110; Photon Systems Instruments, Drasov, Czech Republic). Specifically, the NDVI was calculated as the normalised difference of reflectance at two bands: infrared and red (NDVI = [NIR-R]/[NIR+R]).

### Soil characterisation

Soil physicochemical characterisation was performed according to standardised protocols of the Ionomic Service (Research Support Service, CEBAS-CSIC, Murcia, Spain). The main soil physicochemical characteristics (i.e., pH and mineral content of the soils), were determined in a 1:5 (w:v) soil: water suspension as indicated in the European Standards (EN 13037 1999; EN 13038 1999; EN 13652 2001, respectively), while ammonium was extracted with KCl 2 M (1:5, w:v). In detail, the soil organic carbon (i.e. org. C; pre-treated with HCl to remove the carbonates present in the sample) and total nitrogen (i.e. N) were determined by Elemental Analysis (LECO TruSpec Micro Series, St. Joseph, MI, USA) according to the Dumas combustion method (Buckee 1994). Total potassium (i.e. K) and phosphorus (i.e. P) contents were determined by Inductively Coupled Plasma Optical Emission Spectrometry (ICP-OES) using a Thermo ICAP 6500 Duo equipment (Thermo Fisher Scientific, Waltham, Massachusetts, USA). Results of org. C, N, P or K were expressed as grams per 100 grams of soil (%). Ammonium (NH_4+_) was determined by a colorimetric method based on Berthelot’s reaction (Sommer et al., 1992), while nitrate (NO_3-_), nitrite (NO^2-^) and phosphate (PO_42-_) soil contents were analysed by ion chromatography (HPLC, model 861, Metrohm AG, Herisau, Switzerland). NH_4+_, NO ^-^, NO^2-^, PO ^2-,^ and K results were expressed as µg g^-1^ of soil. The pH was measured using a Crison model 2000 pH meter.

The electrical conductivity (Table 2), an indicator of the concentration of solutes such as cations and anions (Friedman, 2005), was determined for each composite soil sample following the FAO protocol (2021). Specifically, measurements were carried out at room temperature (22°C) using a conductivity meter (HI5321; HANNA Instruments). For each measurement, 20 g of sieved, rock-free soil was mixed with 100 mL of deionised/distilled water (1:5; w/v). The samples were then sealed with bottle caps and placed horizontally in a shaker for 60 minutes under dark conditions. After shaking, the samples were left undisturbed for an additional 30 minutes. The electrical conductivity (expressed as µs cm^-1^) was then measured by carefully inserting the probe of the conductivity meter into the supernatant, ensuring the sediment remained undisturbed.

**Table 2.** Soil characterisation by pH, electrical conductivity, bulk density, water content and nutrients (quantity and stoichiometry) as a function of management practice (i.e. abandoned and managed trees). Means ± SE values (n = 4) are given for each analysed soil variable. Asterisks indicate statistically significant differences between the different types of management (abandoned and managed trees) after one-way ANOVA statistical analysis: * for p < 0.05, ** for p < 0.01, and ns for nonsignificant relationships. When normality and homocedasticity assumptions were not met, the Kruskal-Wallis’s test was used. Abbreviations for soil variables: org. C, soil organic carbon; N, total nitrogen; P, total phosphorus; K, total potassium; N+P+K, is the total sum of nitrogen, phosphorus and potassium.

| Variable | Type of management |  | *p*-value |
| --- | --- | --- | --- |
|  | Abandoned | Managed |  |
| *Physicochemical properties* |  |  |  |
| Soil pH | 3.79 ± 0.12 | 4.19 ± 0.31 | ns |
| Soil Electrical Conductivity (µs cm<sup>-1</sup>) | 24.71 ± 4.32 | 29.99 ± 3.03 | ns |
| Bulk density (g cm<sup>-3</sup>) | 0.52 ± 0.13 | 0.64 ± 0.03 | ns |
| Soil Water Content (%) | 5.48 ± 1.58 | 7.56 ± 4.09 | ns |
| *Soil nutrients* |  |  |  |
| org. C (%) | **8.88 ± 3.15** | **3.19 ± 0.44** | * |
| N (%) | 0.37 ± 0.10 | 0.29 ± 0.03 | ns |
| P (%) | 0.02 ± 0.005 | 0.03 ± 0.001 | ns |
| K (%) | **0.40 ± 0.17** | **1.42 ± 0.09** | * |
| N + P + K (%) | **0.79 ± 0.18** | **1.74 ± 0.10** | ** |
| Nitrate (µg g<sup>-1</sup>) | 26.95 ± 6.24 | 169.66 ± 73.45 | ns |
| Nitrite (µg g<sup>-1</sup>) | 0.71 ± 0.29 | 0.51 ± 0.19 | ns |
| Ammonium (µg g<sup>-1</sup>) | **48.28 ± 15.16** | **10.65 ± 2.23** | * |
| Phosphate (µg g<sup>-1</sup>) | 3.26 ± 0.89 | 2.05 ± 0.36 | ns |
| *Soil stoichiometry* |  |  |  |
| C:N | **23.14 ± 3.59** | **10.75 ± 0.69** | * |
| C:P | **500.34 ± 140.57** | **121.35 ± 20.30** | * |
| N:P | 20.36 ± 3.85 | 11.16 ± 1.42 | ns |

Finally, bulk density (expressed on mass per volume; g cm^-3^) was quantified following the adapted ISO 11272:2017 (Fernandez-Ugalde et al. 2022) and soil water content (%SWC) was quantified as follows:

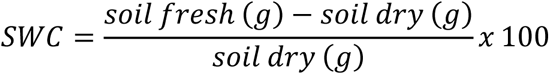

### Assessment of leaf nutrients

Leaf macronutrients (carbon, C; nitrogen, N; phosphorus, P; potassium, K; calcium, Ca; magnesium, Mg; sulfur, S; silicon, Si) and micronutrients (iron, Fe; boron, B; manganese, Mn; sodium, Na; zinc, Zn; copper, Cu; nickel, Ni; molybdenum, Mo) analyses were performed according to standardised protocols of the Ionomic Service (Research Support Service, CEBAS-CSIC, Murcia, Spain). Cations were analysed by ICP-OES using an ICAP series 6500 spectrometer (Thermo Fisher Scientific, Waltham, Massachusetts, USA). Anions were analysed by ion chromatography (HPLC, model 861, Metrohm AG, Herisau, Switzerland) in water extracts obtained by shaking dried leaf material for 2 h. Total C and N content was determined by Elemental Analysis (LECO TruSpec Micro Series, St. Joseph, MI, USA). Results of C, N, P, K, Ca, Mg, S and Si were expressed as % dry weight (g 100 g^-1^) while Fe, B, Mn, Na, Zn, Cu, Ni, Mo were expressed as µg g^-1^ of dry weight.

### Chlorophyll *a* fast fluorescence transient

OJIP was measured with a portable chlorophyll fluorimeter (Fluorpen FP110, PSI, Drásov, Czech Republic) to analyse the photosynthetic apparatus status. This technique estimates the flow of energy through photosystem II (PSII), which is a highly sensitive signature of the linear electron transport flow and the photosynthetic efficiency (Strasser et al. 2000; Stirbet and Govindjee 2011). Excitation via blue light-emitting diodes (455 nm), optically filtered to provide a light intensity of 3000 µmol photons m^-2^ s^-1^, allowed recording of fluorescence transients for 2 s at frequencies of 10 µs, 100 µs, 1ms and 10 ms for time intervals of 10-600 µs, 0.6-14 ms, 14-100 ms and 0.1-2 s, respectively. The fluorescence values at 40 µs (F_o_, step O, all reaction centres of the PSII are open), 100 µs (F_100_), 300 µs (F_300_), 2 ms (step J), 30 ms (step I) and maximal (maximum level of fluorescence, F_M_, step P, closure of all reaction centres) were taken into consideration for further analyses. We further calculated the OJIP parameters (energy fluctuations and yields; more details below) from F_o_ and F_m_ and the fluorescence intensities selected at 40 µs, 100 µs, 300 µs, 2 m, and 30 ms. These parameters are described in Table S2, available as Supplementary data. (i.) The relative variable chlorophyll fluorescence parameters (i.e., V_j_ and V_i_) are derived from steps I and J. (ii.) The specific energy fluxes are expressed per primary quinone acceptor reducing PSII centre: (i.e., ABS/RC, Di_o_/RC, TR_o_/RC, Et_o_/RC). In detail, ABS refers to the photon flux absorbed by the chlorophyll antenna pigments of the PSII. Part of this energy is dissipated, mainly as heat (DI_o_). Another part of this absorbed energy is funnelled to the reaction centre (RC) as trapping flux (TR_o_). In the reaction centre, the excitation energy is converted into redox energy by reducing the electron acceptor QA to QA^-^, which is then reoxidised to QA, creating electron transport (Et_o_). (iii.) Quantum yields and efficiencies (i.e., F_V_/F_M_, Ψo, ϕEo, ϕDo and ϕPav) are directly related to the energetic fluctuations obtained from the specific fluxes per RC (Strasser et al. 2000; Hermans et al. 2003). In detail, F_V_/F_M_, is the maximum quantum yield of primary photochemistry, and represents the probability that an absorbed photon is trapped by the RC and used for primary photochemistry (it is calculated as TR_o_/ABS); Ψo, is the efficiency with which a trapped excitation can move an electron into the electron transport chain further than QA (it is calculated as ET_o_/TR_o_); ϕEo, is the quantum yield of electron transport, representing the probability that an absorbed photon moves an electron into the electron transport chain (it is calculated as Et_o_/ABS); and ϕDo is the quantum yield for energy dissipation. (iv.) Finally, we calculated the performance index (Pi_Abs_), which allows *in vivo* evaluation of plant performance (in terms of biophysical parameters that quantify photosynthetic energy conservation; Strasser et al. 2000). This parameter summarises three important events: light trapping, the quantum efficiency of reduction of QA, and the efficiency of electron transport from quinone to the intersystem carriers of the electron chain (Hermans et al., 2003). Thus, this index is a useful tool to screen photosynthetic performance and characterise plant vitality under stress conditions because it is sensitive to changes in antenna properties, light trapping efficiency, and electron transport (Hermans et al., 2003; Strasser et al., 2000).

### Photosynthetic pigments and tocopherols

The photosynthetic pigments and tocopherols were quantified by liquid chromatography (uHPLC) using the protocol described by Lacalle et al., 2020. Specifically, the extractions were made using 100% pure acetone and the resulting liquid was homogenised (Tissue Tearor model 965670) and then centrifuged at 16,100 g and 4°C. Both were syringe filtered through a 0.22 µm PTFE filter (Whatman, Maidstone, UK). The extractions were performed under cold conditions (4°C) and green light (protecting the samples against direct light). The final quantification of the pigments and tocopherols was made using the ultra-rapid uHPLC method (Lacalle et al., 2020). For this, the extracted samples were injected into an Acquity^TM^ UHPLC H-Class system (Waters®, Milford, MA, USA) using a reverse-phase column (Acquity UPLC® HSS C18 SB, 100Å, 1.8 µm, 2.1 x 100 mm) and a Vanguard^TM^ pre-column (Acquity UPLC HSS C18 SB, 1.8 µm). The photodiode detector (Acquity PDA UHPLC; Waters) was then used to detect pigments in the range of 250-700 nm, while the tocopherols were detected by fluorescence (FLR; UHPLC Acquity, Waters), with an excitation λ of 295nm and an emission of 340 nm. Pigment peaks were integrated at 445 nm. The retention times and conversion factors for carotenoids were the same as those described by (Lacalle et al., 2020). In figures, the following compounds are shown: the total chlorophyll pool, as the sum of chlorophyll a (Chla) and chlorophyll b (Chlb), VAZ, as the sum of V, A and Z, the total carotenoids (t-Car), as the sum of VAZ, lutein, lutein epoxide, α-carotene and β-carotene, α-tocopherol (α-Toc) and the total tocopherols (t-Toc), as the sum of the tocopherol isomers (i.e., δ+ β-γ + α). The VAZ and t-Toc pools were combined into a single variable to simplify the analysis (herein referred to as photoprotective isoprenoids). Chl a + Chl b was expressed on a fresh weight basis (nmol g^−1^ FW). Additionally, the rest of the compounds were expressed on a chlorophyll basis (mmol mol^-1^ Chl).

### Phytohormones profile

The stress hormones ABA, SA, JA, the precursor of ethylene biosynthesis 1-aminocyclopropane-1-carboxylic acid (ACC), OPDA, and the conjugated active form JA-Ile, plus the growth-hormones indole-3-acetic acid (IAA), gibberellins (GA_1_, GA_3_, GA_4_, GA_7_), isopentenyladenosine (IPA), 2-isopentent adenine (2-iP), zeatin (Z) and zeatin riboside (ZR), as well as other hormones like Mel, were all measured using the same methanolic extract via liquid chromatography coupled to electrospray ionisation tandem mass spectrometry (UHPLC/ESI-MS/MS) (Müller & Munné-Bosch, 2011).

In short, 50 mg of freeze-dried needle samples were ground and extracted with 800 μl methanol:isopropanol: acetic acid, 50:49:1 (v/v/v) containing deuterium-labelled internal standards (d6-2-iP, d6-ABA, d4-ACC, d2-GA_1_, d2-GA_4_, d2-GA_7_, d5-IAA, d6-IPA, d5-JA, d4-Mel, d4-SA and d5-tZ) using vortex and ultrasonication (Branson 2510 ultrasonic cleaner, Bransonic, Banbury, CT, United States). Extracts were centrifuged for 10 min at 4°C and 15,980 g. The supernatant was collected, and the entire procedure was repeated twice for a re-extraction of the pellet. Finally, supernatants were mixed and filtered with hydrophobic PTFE filters of 0.22 µm (Phenomenex, Torrance, CA, United States) before analysis and injected into the UHPLC/ESI-MS/MS. Quantification of each compound was performed considering recovery rates for each sample and calibration curves for each analyte.

### Statistical analyses and data processing

One-way ANOVA analyses were performed to look for differences between abandoned and managed *P. radiata* individuals considering the following variables: ecosystem and tree traits (relative irradiance, tree diversity, shrub diversity, herbaceous diversity, fern diversity, LMA, F_V_/F_M_, NDVI, DBH), soil and leaf nutrients, OJIP parameters (V_i_, Vj, F_V_/F_M_, Ψo, ϕEo, ϕDo, ϕPav, Pi_Abs_, ABS/RC, TRo/RC, ETo/RC, DIo/RC), photosynthetic pigments (Chla + Chl b, t-Car, VAZ), tocopherols (α-Toc, t-Toc), the photoprotective isoprenoid pool (VAZ + t-Toc) and phytohormones (ABA, OPDA, SA, ACC, JA, JA-Ile, Mel, IAA, IPA, GA_1_, GA_3_, GA_4_, GA_7_). Before performing one-way ANOVA, we checked the normality (Kolmogorov–Smirnov test), and homoscedasticity (Levene’s test) of all response variables. When these assumptions were not met, the Kruskal-Wallis test was used.

Moreover, a radar plot was performed to determine the main OJIP parameters. For this, we calculated the average values of the OJIP parameters for each type of stand and we standardised them using the abandoned group, for which we used a value of 1 as a reference. Deviations from the 1 value denote an effect in each of the OJIP parameters due to the management stages and ages.

Subsequently, to reduce the dimensionality of the 16 leaf nutrient variables and the 6 stress hormone variables while preserving most of their variation, we performed Principal Component Analysis (PCA). For this, we used the “prcomp” function available in the R-based learning functions. The data was normalised using the scale() function. Based on the obtained leaf nutrient results, the first principal component (PC1) explained 25.13% of the total variance, while the second principal component (PC2) accounted for 16.63%. Likewise, based on the obtained stress-hormone results, the PC1 explained 43.18% of the total variance, while the PC2 accounted for 21.37%. In both cases, the PC1 was used in further analysis as the leaf nutrient status variable and the stress-hormone profile variable, respectively. We also performed Spearman correlations using the “rcorr” function from the “Hmisc” R package (Harrel 2019) to determine the relationship between soil NPK, leaf nutrient status (i.e., PC1, PC2), stress-hormone profile (i.e., PC1, PC2), Mel, photoprotective compounds (i.e., t-Car, VAZ, and t-Toc) and OJIP parameters (F_V_/F_M_ and Pi_Abs_) when considering the abandoned and managed stands. Moreover, to test our hypothesis and based on the above-mentioned Spearman correlation results, we performed multiple sets of linear mixed effect models (LMEs) for each group of study using the “lme” function. The fixed effects of the sets of LMEs included the response and the explanatory variables (i.e. soil NPK, leaf nutrient status (i.e., PC1), stress-hormone profile (i.e., PC1), Mel, t-Car, VAZ, t-Toc and PiAbs), while for the random effect we used “centroid”. Note that no LMEs were run considering F_V_/F_M_ since this parameter strongly covariates with Pi_Abs_ in both management stages. The latter is a more integrative parameter of the photosynthetic performance (Strasser et al., 2000). When normality and homoscedasticity assumptions were not met, we logarithmically transformed the variables to reduce the influence of outliers for model validation. To calculate the coefficients of the LMEs and thus to estimate the effects of each explanatory variable, we ran ANOVAs of either type II (when dealing with a balanced design; i.e., Soil NPK, leaf nutrient status, stress-hormone profile, Mel and Pi_Abs_) or type III (when dealing with an unbalanced design; i.e., t-Car, VAZ, t-Toc) using the car R package; Fox and Weisberg 2019). The residuals of the sets of LMEs were checked for normality using the Kolmogorov–Smirnov test. The final coefficients of the LMEs were estimated using the Restricted Maximum Likelihood Method (REML). The conditional R^2^ values (i.e., the proportion of variance explained by both the fixed and random parts of the LMEs), were calculated using the r.squaredGLMM function (Barton, 2020).

Finally, we run a Structural Equation Model (SEM) based on the results of LMEs. For this, we used the “psem” function (“piecewiseSEM” R package; Lefcheck 2016), which allowed us to introduce random effects (i.e. centroid). We run multiple SEMs to look for all possible causal-effect relationships between soil NPK, leaf nutrient status (i.e., PC1), stress-hormone profile (i.e., PC1), Mel, t-Car, photoprotective isoprenoids (VAZ, t-Toc), and physiological performance (Pi_Abs_). The best models were selected based on stepwise model selection by AIC (i.e., Akaike Information Criterion). Finally, we checked the goodness of fit of the SEM based on Fisheŕs C statistic, which tests whether the model fits the data (when the chi-squared test is *p* > 0.05). Note that all variables were logarithmically transformed before conducting the SEM. All statistical tests were performed using R software (v 4.1.1; R Core Team 2020) in the RStudio platform (RStudio Team 2020).

## Results

### Differences in ecosystem traits, individual traits, needle traits and soil properties between the types of plantations

The diversity of trees, shrubs, and herbs was significantly higher (*p* < 0.05) in abandoned stands than in managed stands. In contrast, no differences were found in the diversity of ferns between stands (Table 1). Despite the greater tree diversity observed in abandoned stands, no differences were found in relative irradiance between abandoned and managed individuals, which means that light availability was approximately equal for all managed and abandoned individuals. We found that managed trees had access to more resources, as a consequence of the reduced diversity in the understorey, compared to abandoned trees.

Needle physiological traits (i.e., LMA and F_V_/F_M_) were found to be significantly higher for managed individuals (with lower DBH) than for abandoned individuals (higher DBH) (*p* < 0.05), although same-aged needles were compared (Table 1). Managed needles had higher LMA (153.42 ± 6.47) while abandoned needles had lower LMA (128.06 ± 5.28). In contrast, F_V_/F_M_ was almost equal in managed needles (0.77 ± 0.004) and in abandoned needles (0.75 ± 0.01). No significant differences in NDVI were observed between treatments.

Soil properties analysis revealed no significant differences in bulk density or soil water content between abandoned and managed stands (Table 2). Besides, no significant differences were found in soil pH and electrical conductivity between the treatments, indicating a strongly acidic soil and low electrical conductivity for both plantations. However, the management practice influenced soil nutrients, org. C content, and soil nutrient stoichiometry ratios (Table 2). In detail, soils in abandoned stands exhibited significantly higher levels of org. C, NH_4+_ and higher ratios of C/N and C/P, while managed-stand soils showed higher values of K and the total NPK.

### Interaction between soil/leaf elemental composition in managed and abandoned stands

Leaf nutrients (i.e. K, Ca, Mg, P, Si) and leaf nutrient stoichiometry (i.e., C/P, N/K, N/P) were affected by management (Figure 2A). Leaves of managed trees showed significantly higher values of Ca (*p*<0.001), Mg (*p*<0.05), P (*p*<0.05), and N/K (*p*<0.001), while leaves of abandoned trees showed significantly higher values of K (*p*<0.01), Si (*p*<0.05), C/P (*p*<0.05) and N/P (*p*<0.05) (Figure 2A). No significant differences in N, C, S and total sum of N+P+K were found between managed and abandoned trees. These results were further supported by the PCA analysis (Figure 2B). While some overlap was observed between groups, samples from the abandoned stands clustered towards more positive values on PC1, whereas samples from the managed stands were grouped towards more negative values on the same axis. This separation highlights the significant influence of management stages on leaf nutrient composition. Specifically, along PC1, micronutrients such as Na, Si, and Cu were the primary contributors, accounting for 61% of the variation. In contrast, PC2 was mainly driven by macronutrients such as P, N, and C, which contributed 79.68% to the observed variance (see Figure S1 available as Supplementary data). By this result, we defined the single variable “leaf nutrient status”, as explained in the Materials and Methods section.

**Figure 2.**
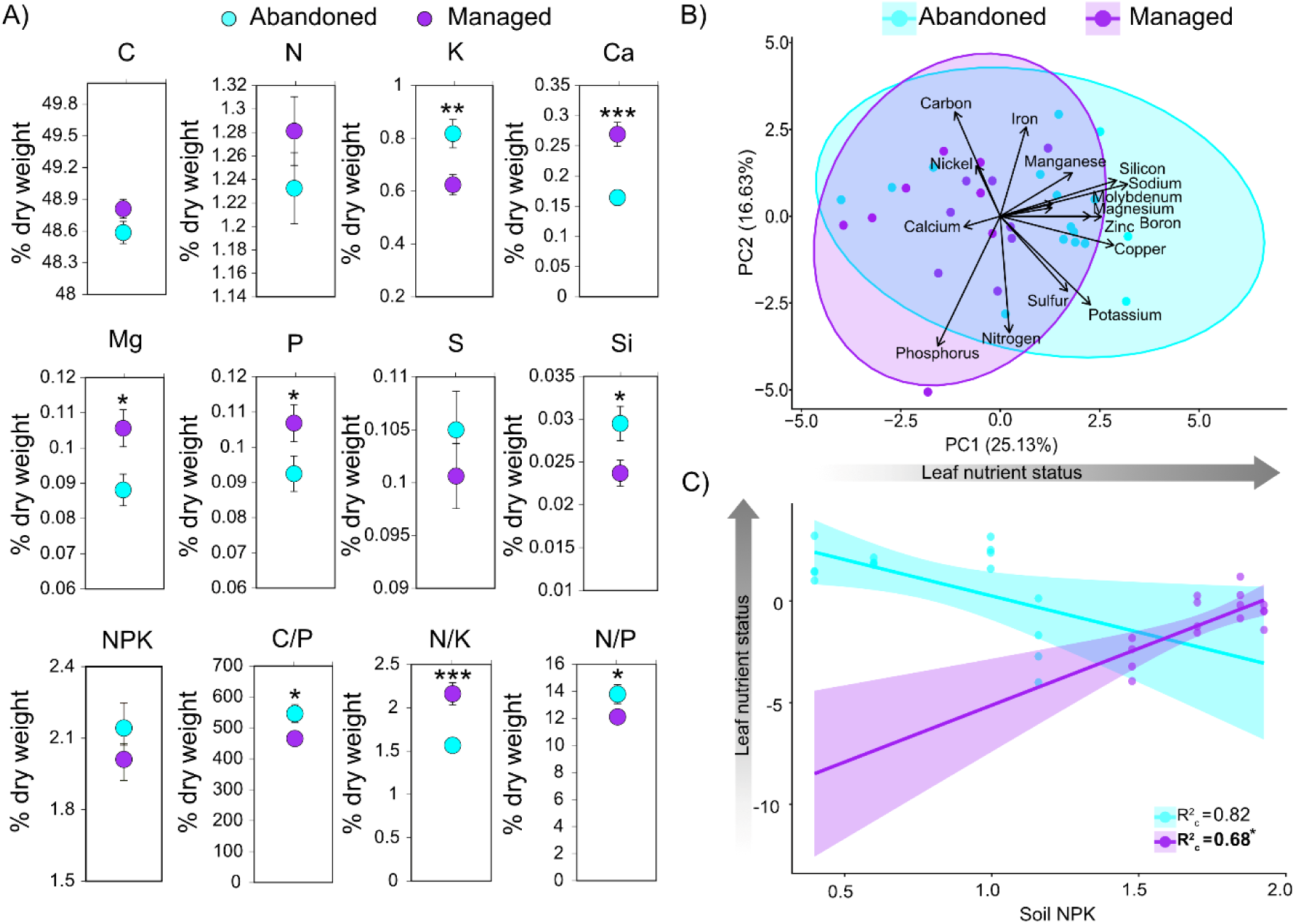
Leaf nutrient analyses of the individual trees in abandoned (blue) and managed (purple) plantations. A) Leaf macronutrient levels expressed as a percentage of dry weight: carbon (C), nitrogen (N), phosphorus (P), potassium (K), calcium (Ca), magnesium (Mg), sulfur (S), silicon (Si), total sum of N+P+K, C/P ratio, N/K ratio and N/P ratio). A one-way ANOVA test was used to compared data between trees growing in managed and abandoned stands. When normality and homocedasticity assumptions were not met, the Kruskal-Wallis test was used. Values are the mean of 16 replicates ± standard error. B) Result of the PCA run to reduce the number of leaf variables and define the single variable “Leaf nutrient status”. The leaf variables considered were the macronutrients carbon, nitrogen, phosphorus, potassium, calcium, magnesium, sulfur and silicon, and the micronutrients iron, boron, manganese, sodium, zinc, copper, nickel and molybdenum. C) Results of the LME model showing leaf nutrient status fitted as a function of soil NPK (% dry weight). Solid lines represent fitted data, while shaded areas represent the 95% confidence interval (abandoned individuals are marked in blue, and managed individuals in purple). Circles represent the different types of management: abandoned individuals (marked in blue) and managed individuals (marked in purple). In the LME, we show the R^2^_c_, measuring the fraction of variation explained by both the fixed and random parts of the models. Bold letters represent significant impact (*p*<0.05) and asterisks indicate the level of statistical significance of the independent variable on the dependent variable (* for *p*< 0.05, ** for *p*< 0.01, and *** for *p*< 0.001).

The effect of soil NPK on the leaf nutrient status within the two forestry management stages (Figure 2C) indicated that in managed trees, high NPK soil concentrations increased the leaf nutrient status significantly (*p*<0.05), indicating dependency of leaf nutrient status on the soil nutrients. On the other hand, no relationship was found in abandoned trees.

### Differences in leaf photoprotective mechanisms and phytohormones between forest stands

Regarding the status of the photosynthetic apparatus, managed trees differed significantly from the abandoned trees in (i) the DI_o_/RC, (ii) the ABS/RC, (iii) and the ϕDo quantum yield (*p* < 0.05; Figure 3A-3B). Indeed, abandoned trees showed higher DI_o_/RC and higher ABS/RC than managed trees. However, no significant differences were found between the managed and abandoned trees regarding the rest of the fluorescence parameters, including F_V_/F_M_ and Pi_Abs_ (Figure 3A and 3B).

**Figure 3.**
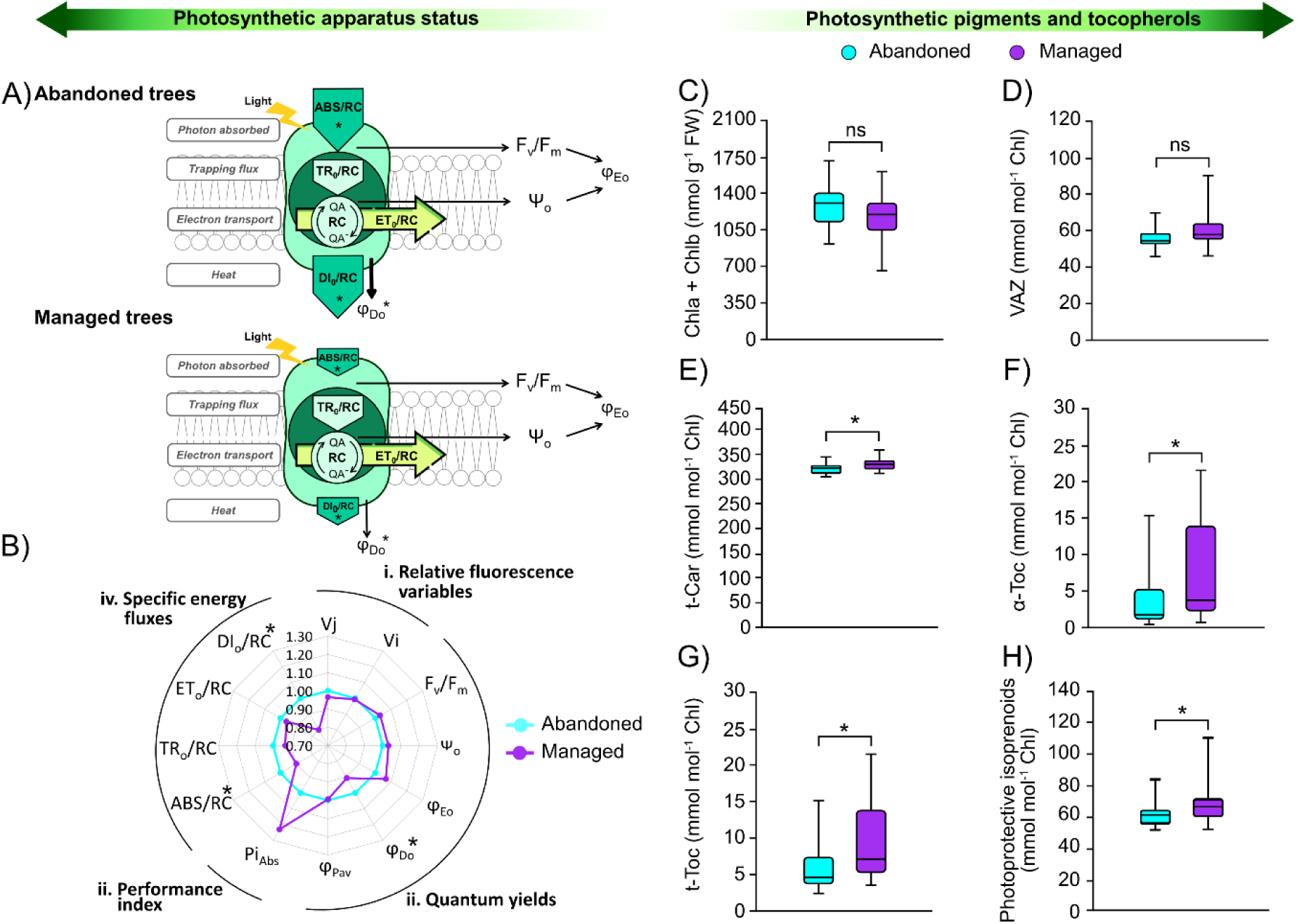
Photosynthetic apparatus status and photosynthetic pigments and tocopherols of abandoned (blue) and managed (purple) pine stands (*n* = 16). A) Simplified representation of the model of energy transfer from solar light to photosynthetic electron transport in the photosynthetic apparatus (adapted from Strasser et al. 2000; Hermans et al. 2003; Encinas-Valero et al. 2022) for abandoned and managed individuals. The size of the triangles and arrows is directly proportional to the change. B) Radar plot depicting the main chlorophyll *a* fluorescence induction parameters derived from the OJIP test where: (i.) the relative chlorophyll fluorescence variable is represented as Vj and Vi; (ii.) quantum yields are represented as F_V_/F_M_, Ψo, ϕEo, ϕDo and ϕPav; (iii.) the potential performance index for energy conservation is Pi_Abs_: and (iv.) the specific energy fluxes per primary quinone acceptor reducing PSII centre are ABS/RC, TRo/RC, ETo/RC and DIo/RC. C-H) Boxplots showing total chlorophylls (Chla + Chlb expressed on a leaf fresh-weight basis, nmol g^-1^ FW; C), the total xanthophyll pool (VAZ expressed on total Chls, mmol mol^-1^ Chl; D), total carotenoids (t-Car expressed on total Chls, mmol mol^-1^ Chl; E), total α-tocopherols (α-Toc expressed on total Chls, mmol mol^-1^ Chl; F), total tocopherol (t-Toc expressed on total Chls, mmol mol^-1^ Chl; G) and total photoprotective isoprenoids (VAZ + t-Toc expressed on total Chls, mmol mol^-1^ Chl; H). Each box represents 50% of the data distribution between the first and third quartile; the central line represents the median, and the upper and lower whiskers cover the 1.5 interquartile range. Asterisks represent significant differences (*p* < 0.05) among abandoned and managed trees. Definitions and formulas for all variables are given in the Materials and Methods, and Table S2 is available as Supplementary data. A one-way ANOVA test was used to compare data between trees growing in managed and abandoned stands. When normality and homoscedasticity assumptions were not met, the Kruskal-Wallis test was used.

The content of isoprenoids (i.e. chlorophylls, carotenoids, and tocopherols) was also analysed in the two forest stands (Figure 3C-3H). Managed and abandoned trees showed no significant differences in Chla + Chlb content (Figure 3C) and in VAZ pool (Figure 3D). However, the content of t-Car (Figure 3E), α-Toc (Figure 3F), the t-Toc pool (Figure 3G) and the total photoprotective isoprenoids (VAZ+t-Toc; Figure 3H) were significantly higher in trees from managed stands than in trees from abandoned stands.

According to the phytohormone PCA (Fig. 4A), a partial overlap was observed between abandoned and managed trees, indicating that both groups exhibited no differences in their overall stress-hormone profiles. This observation is further supported by the complementary analysis shown in Table S1, available as Supplementary data. Specifically, along PC1, ABA contributed the most to the variation (30.73%), followed by OPDA with a contribution of 21.12%. On the other hand, along PC2, SA was the primary contributor (33.94%), followed by ACC (24.87%), the precursor of ethylene (Figure 4B). Based on these findings, the combined variable “stress-hormone profile” was defined as described in the Materials and Methods.

**Figure 4.**
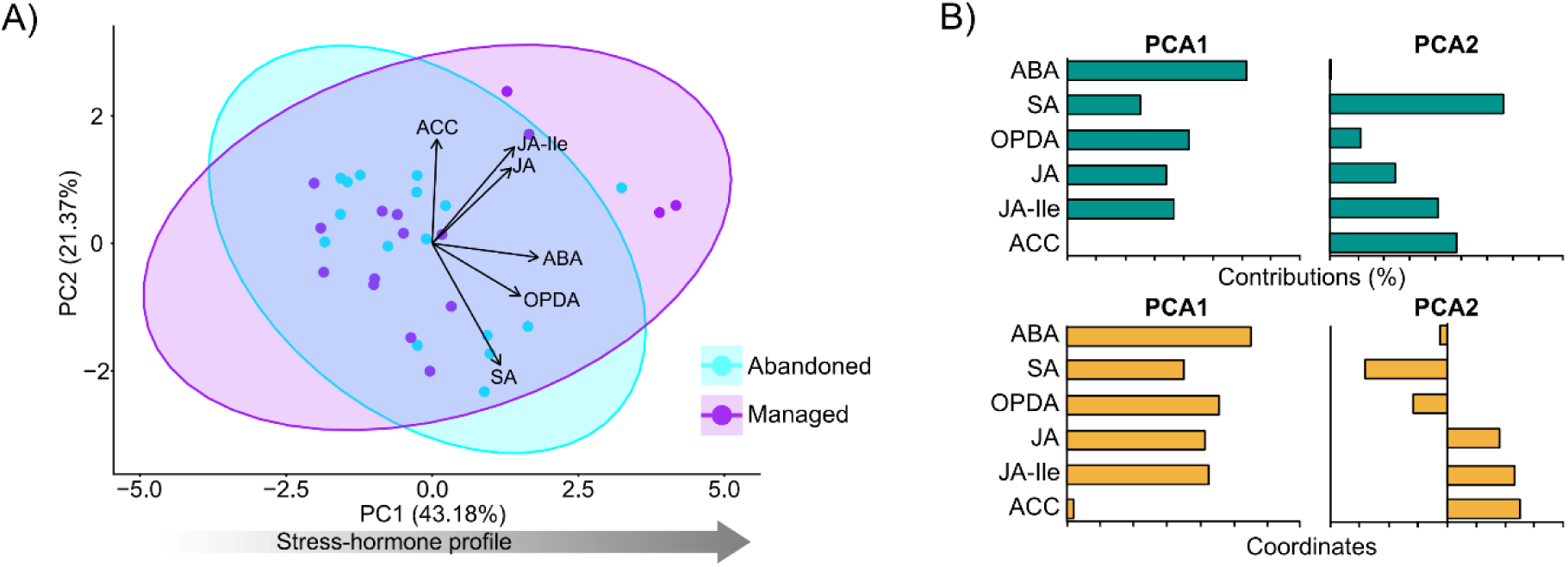
Leaf stress-hormone analyses of individual trees in abandoned (blue) and managed (purple) stands. A) Results of the PCA run to reduce the number of stress hormones and define a single variable (i.e. Stress-hormone profile). B) Bar plots for the contributions (expressed as percentages; green) and coordinates (orange) for each stress-hormone variable in the PCA. The stress-hormone variables considered were abscisic acid (ABA), salicylic acid (SA), the precursor of ethylene 1-aminocyclopropane-1-carboxylic acid (ACC), jasmonic acid (JA) and its precursor 12-*oxo*-phytodienoic acid (OPDA), and the conjugated active form jasmonoyl-isoleucine (JA-Ile) (expressed on a dry weight basis, ng g^-1^ DW).

### Relationships between nutrients, phytohormones, and physiological performance

The Spearman correlation matrix (Figure 5) revealed that variables related to soil nutrients, leaf nutrients, phytohormones (i.e., stress hormones, Mel), and physiological performance (t-Car, VAZ, t-Toc, F_V_/F_M_, Pi_Abs_) significantly correlated in managed and abandoned trees. Specifically, in managed trees, soil NPK levels were correlated positively with leaf nutrient status (PC1; r = 0.56, *p* < 0.05), and negatively with the stress-hormone profiles (PC1; r = -0.74, *p* < 0.01, and PC2; r = -0.52, *p* < 0.05), t-Car (r = -0.59, *p* < 0.05) and VAZ (r = -0.53, *p* < 0.05). Further, leaf nutrient status (PC1) was negatively associated with the stress-hormone profile (PC1; r = -0.63, *p* < 0.01) and t-Car (r = -0.60, *p*< 0.05), while leaf nutrient status (PC2) was negatively correlated with VAZ (r = -0.59, *p*< 0.05). Interestingly, t-Car and VAZ were positively correlated with the stress-hormone profiles PC1 (r = 0.59, *p*< 0.05) and PC2 (r = 0.59, *p*< 0.05), respectively, and t-Toc showed a positive correlation with Pi_Abs_ (r = 0.59, *p*< 0.05). Conversely, no significant correlations were observed between soil NPK and the rest of the variables in abandoned trees. Moreover, leaf nutrient status (PC1) was negatively correlated with the stress-hormone profile (PC1; r = -0.54, *p*< 0.05). Additionally, this stress-hormone profile (PC1) was positively correlated with t-Car (r = 0.60, *p*< 0.05) and t-Car exhibited a negative correlation with Pi_Abs_ (r = -0.63, *p*< 0.01).

**Figure 5.**
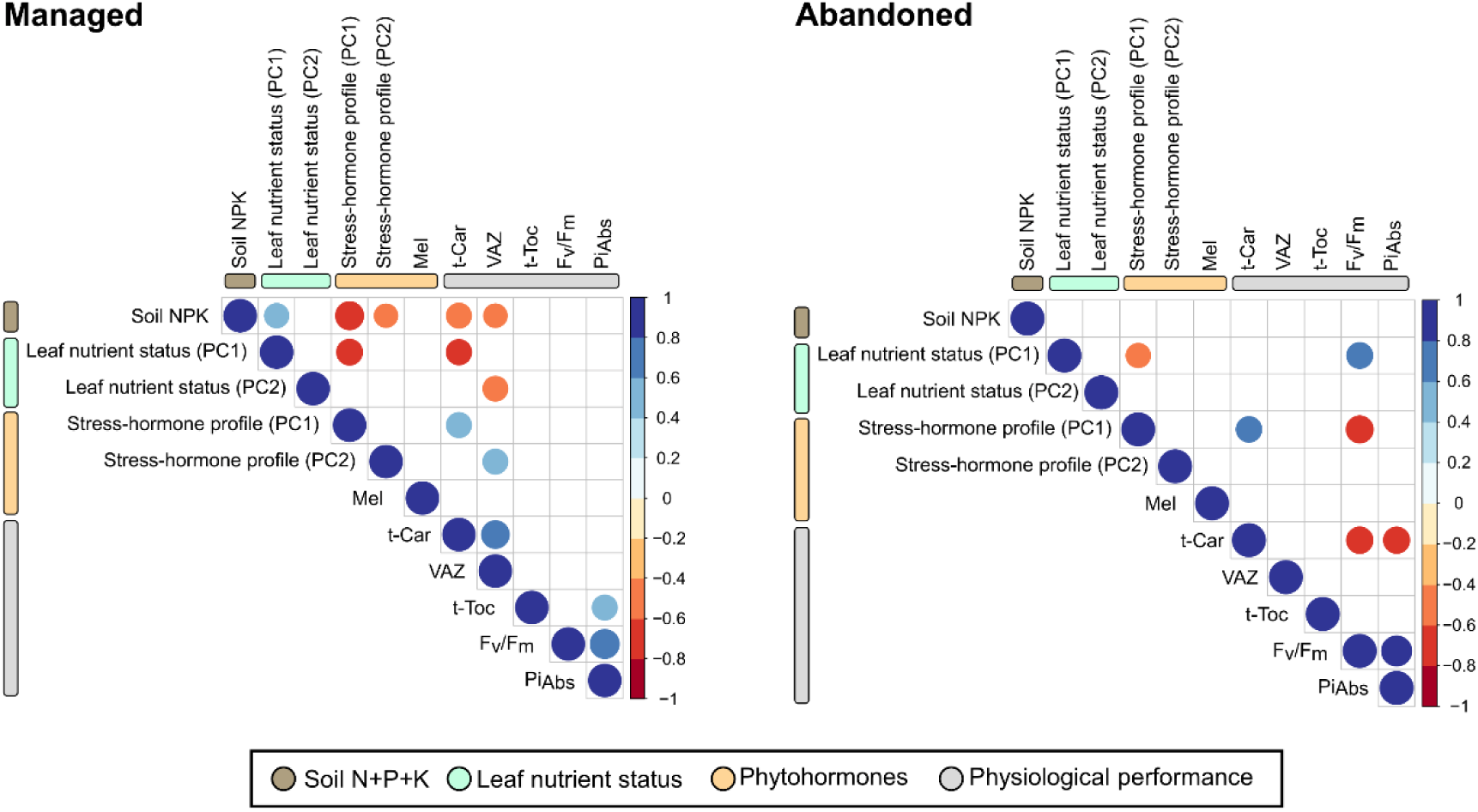
Spearmańs rank correlation matrix plots showing the relationship between: (i.) soil N+P+K (% dry weight); (ii.) leaf nutrient status (i.e., PC1, PC2); (iii.) the stress-hormone profile (i.e., PC1, PC2) and Mel (ng/g DW) both representing phytohormones; (iv.) physiological performance represented by pigments: total chlorophyll pool (Chl a + b, nmol g^-1^ FW), a to b chlorophyll ratio (Chl a/b, mol mol^-1^), total xantophyll pool (VAZ, mmol mol^-1^ Chl), total carotenoid pool (t-Car, mmol mol^-1^ Chl); (v.) total tocopherols (t-Toc, mmol mol^-1^ Chl); and (vi.) chlorophyll *a* fluorescence induction parameters derived from the OJIP test (photochemical efficiency (F_V_/F_M_) and potential performance index for energy conservation (Pi_Abs_). Negative and positive correlations are indicated in red and blue, respectively. The strength of the correlation is indicated by dot size and colour saturation. Only significant correlations are shown (*p*< 0.05).

Based on Spearmańs correlation results, LME analysis (Figure 6) was used to examine the relationship of the following factors with physiological variables: i) soil NPK (Figure 6A-B); ii) leaf nutrient status (Figure 6C-D); iii) phytohormones including stress hormones (Figure 6E-F) and Mel (Figure 6G-H); and iv) physiological performance (Figure 6I-J). In detail, high soil NPK concentrations in managed trees significantly reduced the stress hormone profile (PC1; *p* < 0.001; Figure 6A). Besides, the LME results revealed a significant negative effect of soil NPK on Mel (*P*<0.05; Figure 6B). Neither of these models were significant for abandoned trees (Figure 6A and B). Regarding the models between leaf nutrient status and other variables, managed and abandoned trees showed that both the stress hormone profile (Figure 6C) and t-Car (Figure 6D) had a significant dependency on leaf nutrient status, with a decrease in both variables when leaf nutrient status increased. However, only abandoned individuals showed a significant increase in t-Car when the stress-hormone profile increased (*p*<0.001; Figure 6E). Additionally, a significant negative relationship was found between Mel and the stress-hormone profile in abandoned individuals (*p*<0.05; Figure 6F), while in managed trees an increase in Mel led to an enhancement in leaf nutrient status (*p*<0.05; Figure 6G) and photosynthetic performance (*p*<0.05; Figure 6H). Finally, in managed trees, a significant increase in Pi_Abs_ was observed when photoprotective compounds such as t-Car (*p*<0.05; Figure 6I) and t-Toc (*p*<0.05; Figure 6J) were enhanced. However, in abandoned trees, a significant decrease in Pi_Abs_ occurred when t-Car (*p*<0.05; Figure 6I) increased, thus indicating a differential response due to the management practice.

**Figure 6.**
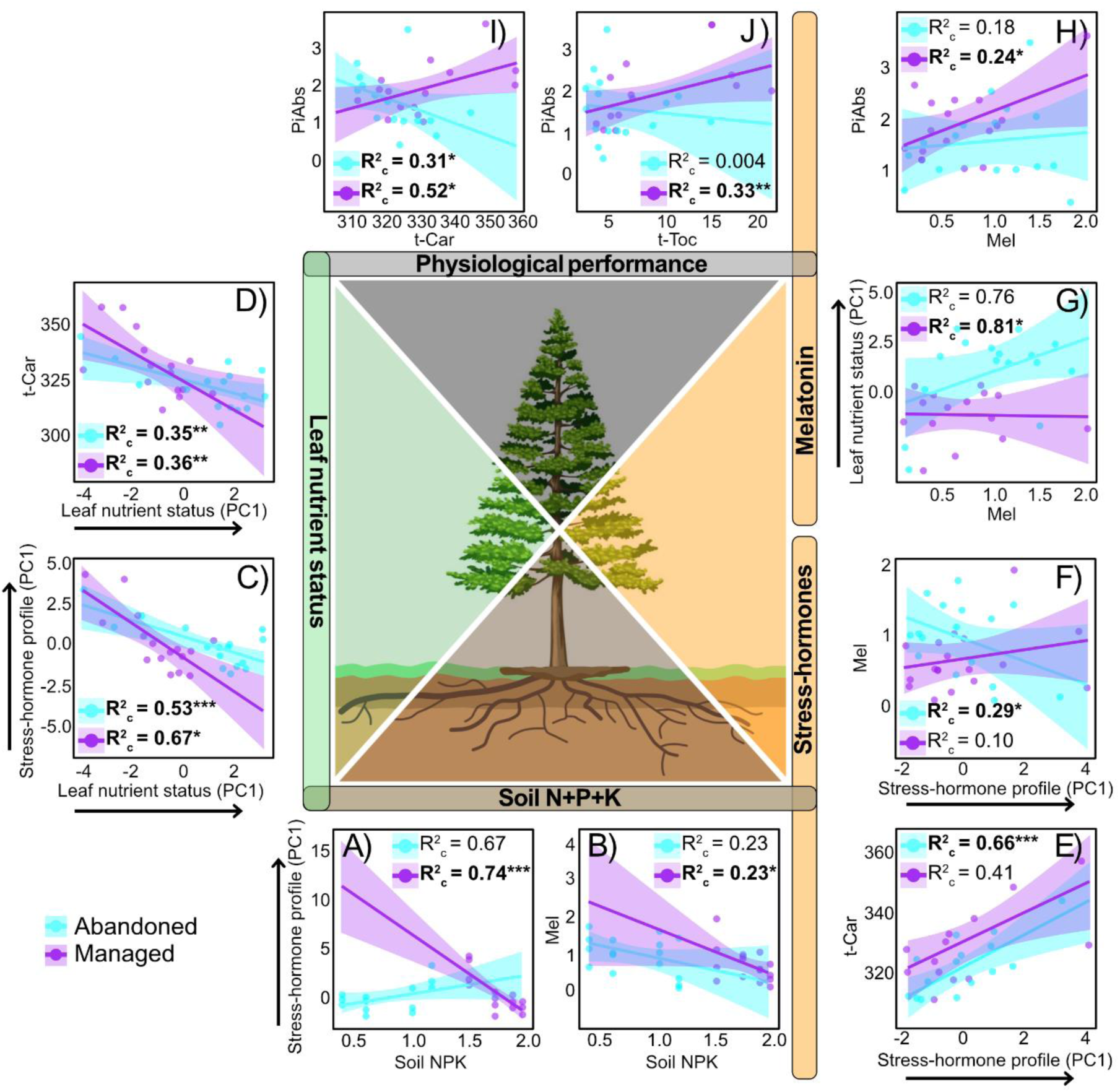
Results of LME models showing the following. i) Plant physiological variables fitted as a function of soil NPK (% dry weight). These physiological variables are stress-hormone profile (A), and Mel (expressed on leaf dry weight basis, ng g^-1^ DW; B). ii) Plant physiological variables fitted as a function of leaf nutrient status. These physiological variables are stress-hormone profile (C) and t-Car (expressed on total Chls, mmol mol^-1^ Chl, D). iii) Plant physiological variables fitted as a function of the stress-hormone profile. These physiological variables are t-Car (expressed on a total chlorophyll basis, mmol/mol Chl, E) and Mel (expressed on a leaf dry weight basis, ng g^-1^ DW; F). iv) Plant physiological variables fitted as a function of Mel. These physiological variables are leaf nutrient status (G) and Pi_Abs_ (i.e., potential performance index for energy conservation; H). v) Pi_Abs_ fitted as a function of photoprotective compounds t-Car (expressed on a total chlorophyll basis, mmol mol^-1^ Chl; I) and t-Toc (expressed on a total chlorophyll basis, mmol mol^-1^ Chl; J). The leaf nutrient status and stress-hormone profile variables were derived from axis 1 of the respective PCAs (Figure 2B and Figure 4A). Solid lines represent fitted data while shaded areas represent the 95% confidence interval (abandoned individuals are marked in blue, and managed individuals in purple). Circles in each plot represent the different types of management: abandoned individuals (marked in blue) and managed individuals (marked in purple). We show the R^2^_c_ for each variable, measuring the fraction of variation explained by both the fixed and random parts of the models. Bold letters represent significant impact (*p*<0.05), and asterisks indicate the level of statistical significance of the independent variable on the dependent variable (* for *p*< 0.05, ** for *p*< 0.01, and *** for *p*< 0.001).

### Distinct causal pathways among nutrients and phytohormones for comparable physiological performance under both forest management stages

The SEM evaluated the causal pathways among soil nutrients, leaf nutrient status, phytohormones (i.e., stress-hormone profile, Mel), and physiological performance (i.e., t-Car, VAZ + t-Toc grouped in photoprotective isoprenoids, Pi_Abs_) (Figure 7). The proposed SEM demonstrated a good fit, as evidenced by Fisher’s C statistics and associated p-value. The abandoned model’s fit was acceptable (ꭓ² = 17.49, p = 0.23), while the managed model exhibited an excellent fit (ꭓ² = 11.28, p = 0.97).

**Figure 7.**
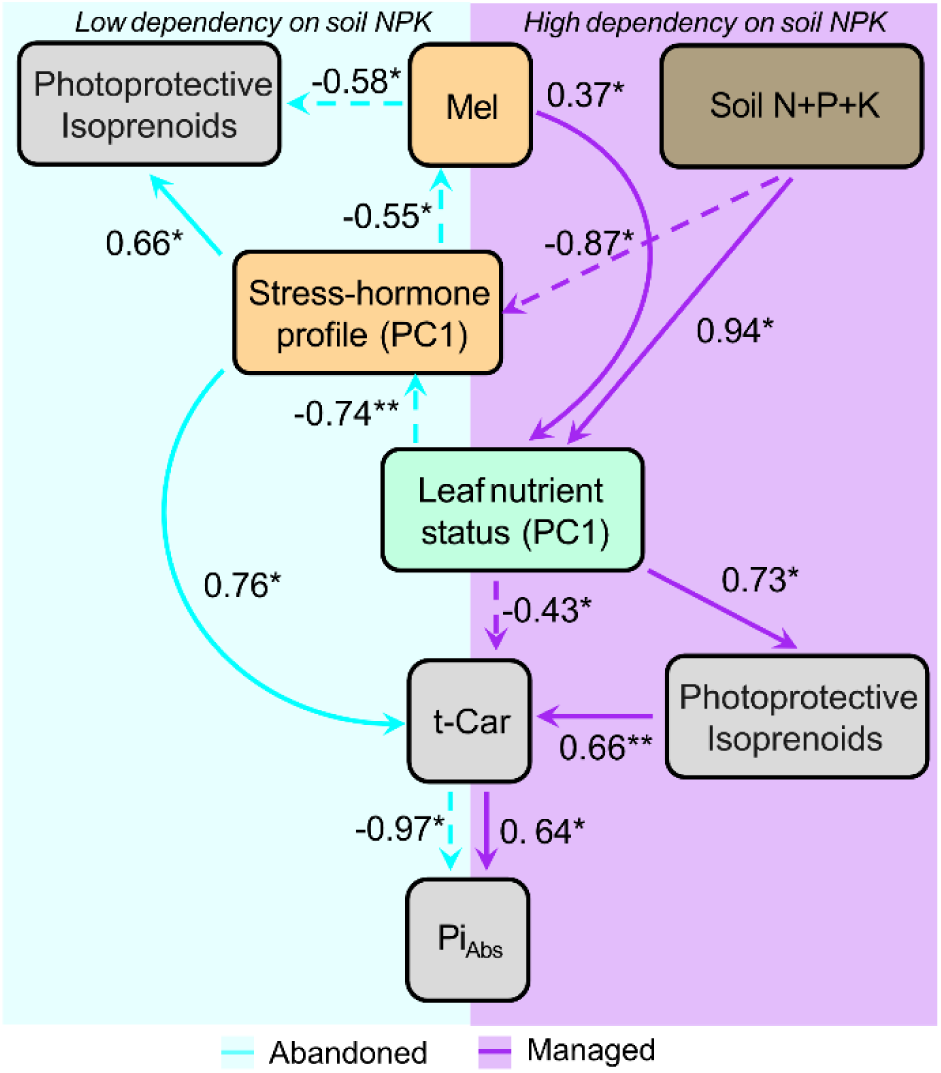
Linkages among nutrients, phytohormones and photoprotective mechanisms in managed and abandoned stands assessed by structural equation modelling (SEM). The path diagram shows a hypothesised causal-effect relationship between soil NPK (N + P + K; %), leaf nutrient status, stress-hormone profile, Mel, photoprotective compounds (t-Car and other photoprotective isoprenoids including VAZ and t-Toc, mmol mol^-1^ Chl) and the potential performance index for energy conservation (Pi_Abs_). The leaf nutrient status stress-hormone profile variables were derived from axis 1 of the respective PCAs (Figure 2B and Figure 4A). All the relationships represented are significant. Arrows indicate causal relationships (positive and negative effects are indicated by solid and dashed lines, respectively); numbers indicate the standardised estimated regression weights, and asterisks represent significant relationships (\**P*< 0.05; \*\**P*<0.01). Blue arrows and shaded areas represent the path diagram of abandoned individuals, and purple arrows and shaded areas represent the path diagram of managed individuals.

Key results of the proposed SEM were as follows. i) Both management stages with differential dependency on soil NPK (low and high for abandoned and managed, respectively) achieved similar Pi_Abs_, however, this was via different pathways. In detail, ii) soil NPK content affected leaf functioning negatively in managed trees, such as leaf nutrient status (decreasing) and the stress hormone profile (increasing), whereas, in abandoned trees, these relationships were not significant. iii) We found a differential role of Mel between the management stages. In managed trees, Mel contributed alongside soil NPK content to the enhancement of leaf nutrient status, but in abandoned trees, Mel and stress hormones influenced photoprotective isoprenoid levels (VAZ and t-Toc). iv) Leaf nutrient status affected t-Car and photoprotective isoprenoid levels in both abandoned and managed stands. In abandoned stands, this influence occurred indirectly through the stress-hormone profile, while in managed stands, it ensued directly. v) Finally, in both cases, t-Car affected Pi_Abs_, but with opposite relationships.

## Discussion

During the summer of 2022, European forests experienced severe dry conditions in the soil (Copernicus Climate Change Service 2022) and increased atmospheric dryness (Gharun et al., 2024). Similar conditions were found for the studied area as indicated by the negative SPEI_2022_ value (–1.33 ± 0.88; Global SPEI database). Soil results (Table 2) indicated that trees in both plantations were growing under nutrient-scarce conditions, as evidenced by low bulk density (Panagos et al., 2024) and low electrical conductivity (Heiniger et al., 2003), both indicative of low nutrient availability, despite managed soils exhibiting significantly higher NPK content than abandoned soils (Table 2). The combination of severe drought and nutrient depletion could compromise leaf-level photosynthetic capacity by reducing the efficiency of the photosynthetic apparatus (Merino et al., 2004). Despite contrasting management stages and soil NPK content (Table 2), both managed and abandoned presented the same chlorophyll content (Fig. 3C) and exhibited a slight downregulation of the photosynthetic apparatus (F_V_/F_M_ close to 0.75 and without significant differences between the two; Table 1). This reduction would be interpreted as a strategy to reduce the risk of excess energy flow towards the production of reactive oxygen species when photosynthesis (carbon assimilation) is restricted under severe drought conditions (Malnoë et al., 2018; Takahashi & Badger, 2011).

However, this similar response of the photosynthetic apparatus masks underlying physiological differences when leaf and soil relationships were considered. We found that managed needles depend on soil nutrient availability for physiological regulation (Fig. 2A, Fig. 6A, B, C, D), with performance declining under nutrient-poor conditions. Whereas abandoned trees maintained independence from nutrient availability (Fig. 2A), displaying long-term acclimation to soil conditions and maintaining photosynthetic function despite low soil nutrient levels, likely through internal nutrient remobilisation mechanisms. These divergent responses reveal a critical trade-off: while management can enhance productivity under nutrient-rich conditions, it may also increase vulnerability to nutrient depletion and environmental stress, particularly under drought. The physiological differences observed, particularly in independence from soil nutrient availability, are likely co-driven by both tree age and management stage. These two factors are inherently confounded in our study, yet together they shape distinct leaf-level physiological strategies. This interaction is crucial for interpreting our findings. Our results highlight that soil nutrient scarcity, especially when combined with drought, can significantly influence tree function by challenging leaf nutrient homeostasis, phytohormones and photosynthetic efficiency. This, in turn, may compromise overall tree resilience. Our study contributes to a deeper understanding of how trees cope with simultaneous nutrient and water limitations, an insight with clear implications for sustainable forest management under climate stress.

### Abandoned needles maintained photosynthetic efficiency while being independent of soil nutrient concentration

In abandoned trees, leaf nutrient status was not influenced by soil NPK availability (represented by PC1; Figure 2C), suggesting that these trees relied less on soil nutrients. Nutrient independence is commonly observed in mature forest stands (Zhang et al., 2024) because mature trees may maximise the nutrients available in the low-fertility soils associated with such mature forests (i.e., low NPK, Table 2), possibly through a mechanism of internal nutrient recycling. This likely involves the translocation of mobile elements, such as P, to meet physiological demands, from older tissues to actively growing tissues without directly relying on soil resources. This is consistent with previous studies that indicated nutrient translocation within plant tissues as a common adaptation in forest plantations established on nutrient-poor soils (Helmisaari, 1995; Laclau et al., 2001; Turner & Lambert, 2014). Elements with higher mobility, such as N, K, and P, are frequently recycled internally, supporting sustained growth and resilience under conditions of limited soil fertility (de Barros Filho et al., 2017). This means that abandoned trees showed stoichiometric flexibility, adjusting their elemental ratios while maintaining constant functions similar to managed trees, such as photosynthetic efficiency (Table 1) and chlorophyll content (Figure 3C; Sistla and Schimel 2012). However, this requires substantial upregulation of the mechanisms of homeostasis and flexibility at the expense of a corresponding energetic cost, e.g. a cost in growth, indicated by the higher N/P ratio, which reflects lower growth rates (Rivas-Ubach et al., 2012), and possibly a reduction in reproductive capacity (Boersma & Elser, 2006).

### Managed trees only maintained photosynthetic capacity with high nutrient content in the leaf

Although both management stages exhibited similar photosynthetic performance, analysis of the photosynthetic apparatus with OJIP curves revealed slight but important differences (Figure 3A-B). Managed trees showed increased light absorbance, as indicated by a higher ABS/RC ratio. This suggested an enhanced ability to utilise light for the synthesis of photosynthates, likely driven by the upregulation of carbon sink capacity (Demmig-Adams et al., 2014). Besides, the increase in Pi_Abs_ associated with Mel in managed trees (Figure 6H) suggests its role in enhancing carbon and nitrogen metabolism (Liang et al., 2018; Ren et al., 2022) as Mel external treatment has been shown to optimise the efficiency of photosystem II and improve CO_2_ assimilation rates (Wang et al., 2019).

Additionally, these trees exhibited higher LMA (Table 1) and increased t-Car, α-Toc and t-Toc contents (Figure 3E-G). These results indicated that managed trees invested more energy per unit leaf area, needing a greater photoprotective demand (Fernández-Marín et al., 2017). Another possibility is that the higher t-Toc content in managed trees may be related to a drought-induced response, potentially indicating a future decline in tree health, which has been reported for other managed species such as holm oaks (Encinas-Valero et al., 2022). Indeed, the latter authors remarked that the upregulation of these isoprenoids represents the final mechanism to protect trees against the combined threats of drought stress and fungal outbreaks, two primary factors driving species decline (Encinas-Valero et al., 2022). However, it is important to note that physiological responses to interacting causal factors like these may vary across different forest ecosystems. Consequently, the upregulation of t-Toc may not act universally as a warning signal for decline (Cavender-Bares & Logan, 2022). Indeed, the elevated photoprotective isoprenoids (Figure 3H) in managed trees may contribute to improved overall performance, as shown by the higher Pi_Abs_ (Figure 6I, J and Figure 7). These responses appear to be driven by both leaf nutrient status and soil nutrient availability (Figure 7), underscoring a nutrient-driven strategy that prioritises photosynthesis and productivity. This strategy is further supported by Mel’s regulatory role in nutrient acquisition under stressful conditions or nutrient deficiencies (Sun et al., 2022). However, this strategy reflects an important trade-off: while managed trees may achieve higher photosynthetic rates than abandoned trees, their vitality could be compromised when leaf nutrient content is low, making these trees more vulnerable to nutrient imbalances or environmental stressors over time.

### Physiological stress from nutrient deficiency affected only managed trees, while, potassium homeostasis supported drought tolerance in abandoned trees

The nutritional status of leaves and trees is closely associated with their performance and survival, especially under environmental stresses like drought (Gessler et al., 2017). The dependence of managed trees on external soil nutrients may imply a higher susceptibility to drought stress because soil nutrient availability is often reduced under drought conditions (Gessler et al., 2017). In managed trees, soil NPK availability (Figure 6A) and leaf nutrient status (Figure 6C) significantly influenced the production of stress hormones (i.e. ABA, jasmonates), suggesting that soil nutrient deficiencies induced physiological stress in these stands. This is consistent with previous studies indicating that nutrient scarcity triggered the synthesis of ABA and other stress hormones, which play critical roles in maintaining homeostasis under resource-limited conditions (Rubio et al., 2009).

Our results also revealed higher K concentrations in abandoned leaves than in managed ones (Figure 2A-B). Potassium is crucial for controlling stomatal regulation (Johnson et al., 2022) and root morphology (Xu et al., 2021), particularly under drought conditions. Besides, in many cases, K-deficient plants tend to be more susceptible to infection than those with an adequate supply of K (Sardans & Peñuelas, 2021). The ability of abandoned trees to maintain K homeostasis may be a physiological advantage in the face of drought, potentially improving their ability to maintain water-use efficiency and then minimising water loss. Overall, the ability of abandoned trees to adjust their elemental composition by internal nutrient cycling mechanisms enabled them to maintain key physiological functions, such as photosynthesis (Figure 3A-B), without a strong reliance on external soil nutrients. This strategy likely contributes to their resilience, particularly in nutrient-poor and dry soils. In contrast, managed trees, while benefiting from higher soil nutrient availability, may be more vulnerable to nutrient depletion and drought (Romanyà & Vallejo, 2004), emphasising the need for careful management strategies to promote long-term forest health and resilience.

### A key role of Mel in natural environments

Mel has been identified in a wide range of plant species across various taxonomic groups, primarily in herbaceous plants and crops (Arnao et al., 2022 and references therein). In this study, we report for the first time the presence of Mel in *P. radiata* growing under natural environmental conditions. Our results also highlight the potential role of Mel in nutrient uptake (Figure 6B-G and Figure 7). In managed trees, Mel improves leaf nutrient status (Figures 6G and 7) under soil nutrient deficiencies (Figure 6B). In contrast, this relationship was not observed in abandoned individuals, where Mel, together with stress hormones, showed a causal relationship with photoprotective isoprenoid concentration (Figures 6F and 7). The distinct relationships observed for Mel in both stages (Figure 6F, G and 7) highlight its versatile functions. In managed trees, Mel may facilitate nutrient acquisition by modulating root architecture(C. Liang et al., 2017) or nutrient transport activity (Shahani et al., 2023), since it has been shown to reorder ion homeostasis and influence overall mineral nutrition (Huang et al., 2022; Sun et al., 2022).

Contrasting the resource strategies highlights the impact of the management stages on physiological processes. Managed systems prioritise overall vitality (physiological performance) while abandoned systems focus on long-term acclimation, emphasising defence and survival. These management-specific physiological traits are crucial for ecosystem processes and functions. In conclusion, our findings showed that the management stages and ages plays a critical role in determining whether trees allocate their resources towards physiological efficiency or resilience, with implications for forest health and ecosystem services.

## Authors’ Contributions

Study conception and design by RE, IA and JCY. Research performance and field sampling by LRL, FSO, AM, UA, CM, UOB, US, LP, JCY and RE. Experiment execution and data collection by LRL, FSO, UA, SMB, TM and UOB. Data analyses by LRL, FSO. Data interpretation by LRL, SMB, JCY and RE. Manuscript writing by LRL and RE. All the authors participated in the final writing and revision.

## Acknowledgements

We thank Ana-Maria Hereş for her help during field sampling and the revision of the manuscript. We also acknowledge the CEBAS-CSIC Ionomic Service (Murcia, Spain) for the nutrient analysis. Thanks to Laurence Cantrill for the English editing.

## Data Availability Statement

The data that support the findings of this study are available from the corresponding author upon reasonable request.

## Supporting Information

Additional supporting information can be found online in the Supporting Information section.

