## Supplementary material for "Contrasting needle physiological strategies to soil nutrient scarcity in radiata pine plantations": Table S1

**Table S1.** Phytohormones in needle tissue (expressed on a dry weight basis, ng g^-1^ DW) of individual trees in abandoned and managed stands. Mean ± SE values (n = 16) are given for each analysed hormone variable. An asterisk indicates the parameters where abandoned and managed individuals are significant (*P* < 0.05).

|  | Type of management | | |
| --- | --- | --- | --- |
| Variable | Abandoned | Managed | *P-*value |
| Stress hormones |  |  |  |
| ABA | 262.93 ± 51.50 | 414.55 ± 126.77 | ns |
| SA | 1725.60 ± 482.62 | 1003.76 ± 240.16 | ns |
| OPDA | 23.74 ± 4.11 | 30.33 ± 5.84 | ns |
| JA | 42.94 ± 9.39 | 80.34 ± 27.74 | ns |
| JA-Ile | 3.87 ± 2.17 | 6.90 ± 3.34 | ns |
| ACC | 131.76 ± 13.34 | 134.88 ± 11.23 | ns |
| Growth hormones |  |  |  |
| IAA | 50.77 ± 7.84 | 36.51 ± 8.45 | ns |
| GA1 | 22.60 ± 5.54 | 23.60 ± 6.02 | ns |
| GA3 | 59.55 ± 5.60 | 83.53 ± 11.47 | ns |
| GA4 | 502.84 ± 18.34 | 468.39 ± 11.93 | ns |
| GA7 | 3.09 ± 0.50 | 2.60 ± 0.22 | ns |
| IPA | 0.98 ± 0.23 | 2.72 ± 1.31 | ns |
| 2-iP | 0.60 ± 0.23 | 0.39 ± 0.14 | ns |
| Z | 0.24 ± 0.07 | 0.39 ± 0.14 | ns |
| ZR | 2.30 ± 0.38 | 6.90 ± 2.18 | ns |
| Other hormones |  |  |  |
| Mel | 0.96 ± 0.13 | 0.68 ± 0.11 | ns |
