## Supplementary material for "Contrasting needle physiological strategies to soil nutrient scarcity in radiata pine plantations": Table S2

**Table S2.** Definitions of terms and formulas for the OJIP-test parameters used in the analysis of the chlorophyll *a* fluorescence transient are shown in Figure 3, based on the equations outlined by Strasser et al. (2000).

| Data extracted from the recorded fluorescence transient OJIP | |
| --- | --- |
| F_t_ | Fluorescence at time t after the onset of actinic illumination |
| F_o_ ≡ F_50 µs_ | Minimal fluorescence intensity at 50 µs, when all reaction centres (RCs) are open |
| F_j_ ≡ F_2 ms_ | Fluorescence value at 2 ms (J-level) |
| F_i_ ≡ F_30 ms_ | Fluorescence value at 30 ms (I-level) |
| F_m_ ≡ F_p_=F_1 s_ | Maximal fluorescence intensity when all RCs are closed |
| M_o_ = 4(F_300 µs_–F_o_)/(F_M_–F_o_) | The initial slope of the fluorescence transient |
| S_m_ Area/(F_m_-F_o_) | Normalised area (assumed proportional to the number of reductions and oxidation of one QA-molecule during the fast OJIP transient, and therefore related to the number of electron carriers per electron transport chain) |
| Fluorescence parameters derived from the extracted data | |
| V_t_ = (F_t_-F_o_)/(F_m_-F_o_) | Relative variable chlorophyll (Chl) fluorescence at time t (from F_o_ to F_m_) |
| V_j_ = (F_j_-F_o_)/(F_m_-F_o_) | Relative variable Chl fluorescence at 2 ms (at the J-step) |
| V_i_ = (F_i_-F_o_)/(F_m_-F_o_) | Relative variable Chl fluorescence at 30 ms (at the I-step) |
| Specific energy fluxes per RC (where TR, ABS, and ET denote the trapped and the absorbed excitation energy fluxes, and the electron transport rate, respectively) | |
| ABS/RC (M_o_/V_j_)^.^F_m_/(F_m_–F_o_) | Specific flux for absorption: absorption flux per RC. Also, a measure of PSII apparent antenna size |
| TR_o_/RC M_o_/V_j_ | Specific flux for trapping: trapped energy flux per RC, resulting in the reduction of QA to QA^–^ |
| ET_o_/RC (M_o_/V_j_)(1–V_j_) | Specific flux for electron transport: electron transport flux per RC |
| DI_o_/RC (M_o_/V_j_)(F_o_/F_v_) | Specific flux for dissipation: the excitation energy dissipated, mainly as heat and less as a fluorescence emission per RC |
| Quantum yields or flux ratio | |
| φ_Po =_ F_V_/F_M =_ TRo/ABS | The maximum quantum yield of primary photochemistry represents the probability that an absorbed photon is trapped by the RC and used for primary photochemistry. |
| Ψ_o =_1–V_j=_ETo/TRo | The efficiency with which a trapped exciton can move an electron into the electron transport chain further than QA. |
| φ_Eo =_ (1–(F_o_/F_m_)) =ETo/ABS | The quantum yield of electron transport: represents the probability that an absorbed photon moves an electron into the electron transport chain |
| φPav = | φ_Po_ (SM / tFm ) tFm = time to reach Fm (in ms) |
| φ_Do=_1– φ_Po_–(F_o_/F_m_) | The quantum yield for energy dissipation |
| Pi_Abs=_[RC/ABS][TR_o_/(ABS-TR_o_)][ET_o_/(TR_o_-ET_o_) | Performance index (potential) for energy conservation from photons absorbed by photosystem II to the reduction of intersystem electron acceptors. |
