## Supplementary material for "Contrasting needle physiological strategies to soil nutrient scarcity in radiata pine plantations": Figure S1

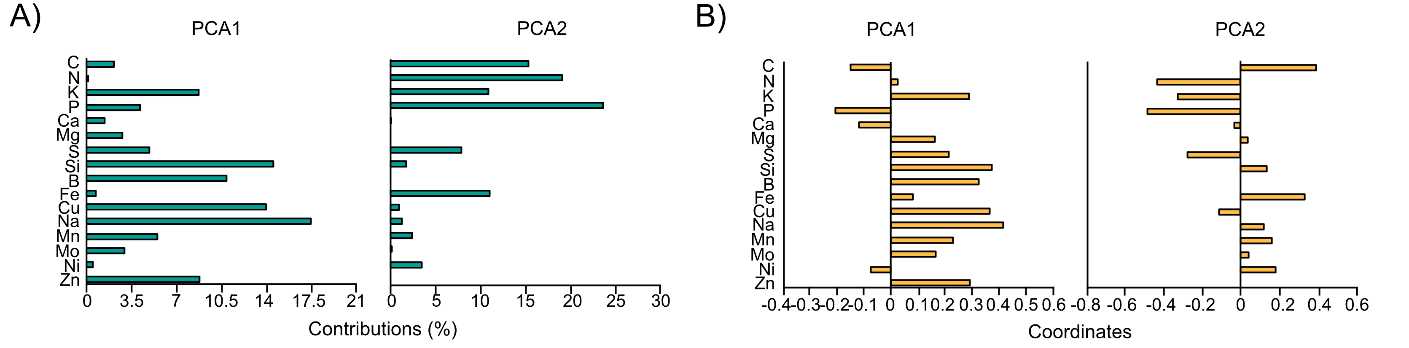

**Figure S1.** Bar plots for the contributions (expressed as percentages; green; A) and coordinates (orange; B) for each leaf nutrient variable in the PCA (Figure 2).
